# Defining the constituents and functions of a lipid-based jumbo phage compartment

**DOI:** 10.1101/2024.03.24.586471

**Authors:** Deepto Mozumdar, Andrea Fossati, Erica Stevenson, Jingwen Guan, Eliza Nieweglowska, Sanjana Rao, David Agard, Danielle L. Swaney, Joseph Bondy-Denomy

**Affiliations:** Department of Immunology and Microbiology, University of California San Francisco, San Francisco 94158 CA, USA; J. David Gladstone Institutes, San Francisco 94158 CA, USA; Quantitative Biosciences Institute (QBI), University of California San Francisco, San Francisco 94158 CA, USA; Department of Cellular and Molecular Pharmacology, University of California San Francisco, San Francisco 94158 CA, USA; Department of Biochemistry and Biophysics, University of California, San Francisco, San Francisco, CA, 94158 USA; Chan Zuckerberg Imaging Institute, Redwood City, CA, 94065 USA

## Abstract

Viruses are vulnerable to cellular defenses at the start of the infection when a single copy genome has not yet initiated *de novo* viral protein synthesis. A family of jumbo phages related to phage ΦKZ, which infects *Pseudomonas aeruginosa,* uses a protein-based phage nucleus to protect the DNA during middle and late stages of infection, but how it is protected prior to phage nucleus assembly is unclear. To reveal the environment surrounding injected phage DNA, we determine which proteins interact with phage proteins injected with the phage genome. Here, we surprisingly identify host proteins related to membrane and lipid biology co-purifying with the injected phage protein. Staining of infected cells with lipophilic dyes revealed a clustering of host lipids co-localized with phage DNA and protein in an early phage infection (EPI) vesicle. Early gene expression is induced by the co-injected virion RNAP (vRNAP), with specific early proteins then associating with the EPI vesicle and ribosomes, likely to facilitate localized translation. As the infection progresses, the phage genome separates from the EPI vesicle and vRNAP, moving into the nascent proteinaceous phage nucleus. Enzymes involved in DNA replication and CRISPR/restriction immune nucleases notably do not localize to the EPI vesicle, supporting that DNA is not exposed at this early time point. We therefore propose that the EPI vesicle is rapidly constructed together with injected phage proteins, phage DNA, host lipids, and host membrane proteins to enable genome protection, early transcription, localized translation, and to ensure the faithful transfer of the single copy phage genome to the proteinaceous nucleus.

## INTRODUCTION

ΦKZ-like jumbo phages infecting *Pseudomonas, Serratia, E. coli, Vibrio,* and *Salmonella* build a large proteinaceous nucleus-like compartment^1,2^, that houses the replicating bacteriophage DNA and selectively excludes diverse DNA-targeting bacterial immune nucleases^3,4^. The compartment itself is constructed of a large protein lattice made up of a single protein called PhuN or Chimallin subunit A (ChmA)^5,6^, encoded as gp54 in phage ΦKZ^2^. This structure is centered in the cell by a tubulin-like protein called PhuZ^7^, where the nucleus often grows larger than the width of the cell during phage DNA replication. The phage encodes DNA polymerase and a non-virion RNA Polymerase (nvRNAP) complex^8–10^ that are presumed to be responsible for replication and transcription, respectively, inside the nucleus-like structure^1^. mRNAs are translated in the cytoplasm while some phage and host proteins are imported into the phage nucleus^1^. Capsid and tail assembly proceeds in the cytoplasm and the phage DNA is loaded at the phage nucleus periphery where capsids dock to receive the phage genome^1,7^. Throughout these latter stages of infection, the phage genome is therefore completely protected from host nucleases^3,4^ and displays remarkable viral organization akin to virus factories assembled by eukaryotic viruses.

Despite generally strong knowledge about the middle and late stages of infection^1–4,7,11^, the early stages remain poorly defined. ΦKZ packages and ejects a virion RNAP (vRNAP) complex^9,12^ along with its genome, which initiates rapid transcription. The major phage nucleus protein (PhuN/ChmA) is rapidly transcribed and translated to assemble and protect the phage genome during replication, but it is not built until ∼20-30 minutes post-infection^1,13^. How the injected genome is protected from immune nucleases and what its surroundings are shortly after injection is currently unknown. We have previously presented two lines of evidence that the phage DNA is protected from nucleases by an entity distinct from the phage nucleus: i) ΦKZ resists endogenous and over-expressed restriction enzymes and CRISPR enzymes that act quickly during infection, suggesting that the injected DNA is not immediately exposed for degradation^3^, ii) upon infection arrest via Cas13a-induced depletion of the transcript encoding PhuN/ChmA, phage DNA was stably present in the cell for >1h despite the presence of an active Type I restriction-modification system^3^. Following these observations, thin-section transmission electron microscopy studies observed that ΦKZ (infecting *P. aeruginosa*) and a related phage SPN3US (infecting *S. typhimurium*) generate small round organelles early in the infection^13,14^. More recent studies have observed that cells infected with 201Φ2-1 phage (*P. chlororaphis*) or Goslar phage (*E. coli*) possess “<u>u</u>nidentified <u>s</u>pherical <u>b</u>odies (USBs)” or “<u>e</u>arly <u>p</u>hage infection (EPI) vesicles”, which were proposed to contain a lipid bilayer based on cryo-electron tomography (cryo-ET)^6,15^. The function and constituents of this early organelle, whether it derives from the host inner membrane, and whether injected phage DNA or protein are inside remains unclear.

To understand the processes that ensure faithful early infection, we sought to identify host and phage proteins that interact with the injected phage genome by tracking co-injected phage proteins. The “inner body” is a large cylindrical proteinaceous structure in the phage head^16–19^ (**Figure 1A**). A previous mass spectrometry (MS) study with tailless virions reported six highly abundant virion proteins (gp89, 90, 93, 93, 95, 97, 162) that likely constitute the majority of this IB structure^19^. Our previous work using unbiased size exclusion chromatography-MS (SEC-MS) identified the IB interactome and immunoblotting demonstrated that 8 proteins are injected with the phage genome^20^. Additionally, fluorescence microscopy revealed that high copy gp93 is also deposited in the cell with the phage genome^21^. Here, we use gp93 and gp95 as bait to identify the host and phage constituents of an early organelle that appears immediately at the start of infection. The injected protein and DNA cargo of ΦKZ-like jumbo phages associates with lipids derived from the bacterial membrane to form a lipidic compartment distinct from the protein-based nucleus, which is a hub for early phage infection.

**Figure 1.**
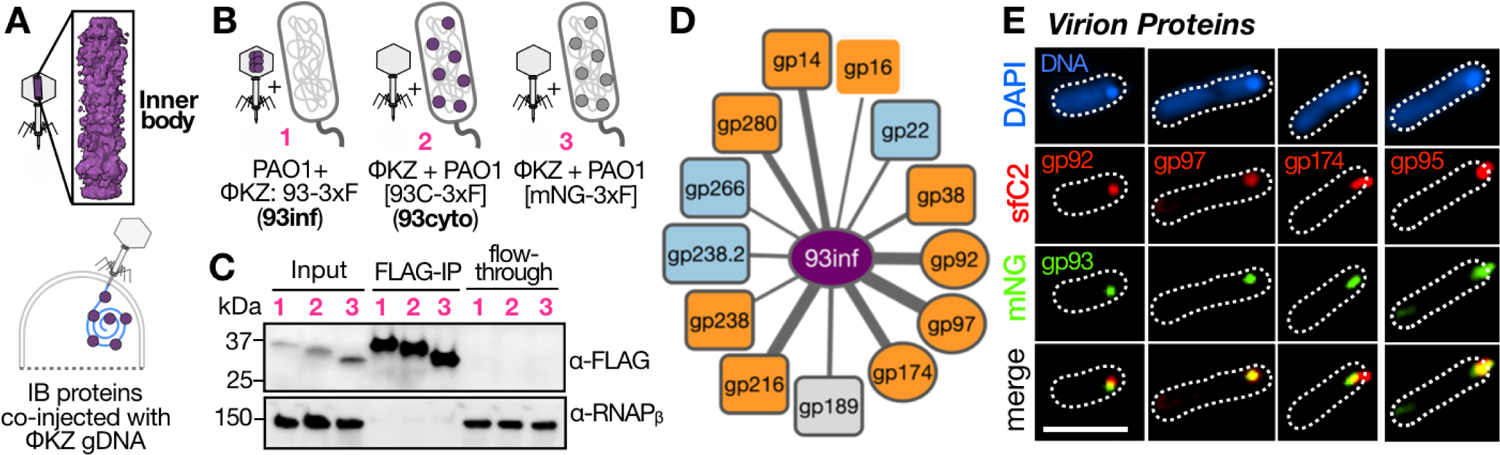
AP-MS with inner body protein gp93-3xFLAG reveals the proteinaceous constituents of a putative early structure. **(A)** During infection, the proteinaceous inner body^18,19^ of ΦKZ is co-injected into bacterial cells with the phage genomic DNA. **(B)** Cartoon schematic illustrating the setup of the FLAG AP-MS experimental samples namely (1) PAO1 cells infected with ΦKZ: gp93-3xFLAG (93inf), PAO1 cells expressing an N-terminal truncated gp93C-3xFLAG infected with ΦKZ (93cyto) or (3) mNeonGreen-3xFLAG infected with ΦKZ. **(C)** Representative ɑ-FLAG western blot of 3xFLAG-tagged protein levels from MS samples in cellular lysate input, FLAG-IP and flow-through. ɑ-RNAPᵦ is used as a loading control. **(D)** Interactome map of ΦKZ proteins that are detected in AP-MS with BFDR<0.05 in 93inf as compared to mNG-3xF and 93cyto negative control. Colors: orange = phage proteins that co-localize with injected gp93-mNG, Blue = phage proteins that do not co-localize with injected gp93-mNG, Grey = microscopy/ validation not attempted; Grey border = that PPI was also found at high confidence (BFDR <= 0.05) with phage injected gp95. Line width is proportional to the average spectral count of the interacting protein (wider width = higher spectral count). Object shape: Rounded rectangle = non-virion protein, Oval = virion protein. **(E)** Representative fluorescence microscopy images of PAO1 cells infected with ΦKZ: gp93-mNeonGreen (mNG; green) packaged with gp92/95/97/174-sfCherry2 (sfC2; red). Cells stained with DAPI to visualize DNA. Scale bars = 5 µM. See also **Supplementary Figure S1** and **Supplementary Table S1**.

## RESULTS

### The injected protein cargo of ΦKZ interacts with host metabolic and membrane proteins

We selected gp93 as a bona fide injected protein^20,21^ to use as bait for AP-MS experiments hoping to reveal the early environment proximal to the phage genome. Fluorescent labeling experiments with other IB proteins revealed that gp90, gp95, and gp97 are also injected with the phage genome, while gp89 and gp162 remain in the head (**Supplementary Figure S1A**). Moreover, bioinformatics analysis of these injected proteins reveal that gp93, gp95, gp97, gp162, and gene neighbors gp92, gp94 and gp163 all share a paralogous domain (PF12699.11) and are proteolyzed during packaging, suggesting that these proteins participate in a shared functionality (**Supplementary Figure S1B, S1C**). Given the high protein copy number in the virion, clear patterns of injection and the previous biochemical observation that gp93 associates with the phage genomic DNA ^19^, we next assessed what gp93 interactors can reveal about the environment the ΦKZ genome finds itself in shortly after the start of infection. A similar experiment was conducted with gp95 and resulting interactors will be compared.

To identify gp93 interactors, gp93 with a C-terminal 3xFLAG affinity tag was packaged into the phage virion to conduct affinity purification during infection (“gp93inf”) and subsequent MS analysis. Interactors that were specifically enriched in the injected gp93 purification were assessed by comparing the detected protein intensities to experiments in which gp93-3xFLAG was expressed in the cytoplasm (i.e. not entering the cell from the phage virion, “gp93cyto”) and to an mNeongreen-3xFlag negative control (Figure 1B). Infections were arrested with gentamicin to limit *de novo* synthesis and synchronize the protein harvest time point. This preferentially captures proteins injected from the phage or recruited from the host, as opposed to proteins synthesized in response to phage infection. Western blotting was used to confirm that the 3xFLAG constructs were expressed and pulled down via FLAG-IP at equivalent levels (Figure 1C). A total of 29 proteins were identified as high-confidence interactors of injected gp93, consisting of 13 phage proteins (**Supplementary Table S1**) and 16 host proteins (**Supplementary Table S2**). gp93 interactors fell into three groups: virion proteins (Figure 1D, oval), non-virion phage proteins (Figure 1D, rounded rectangle), and host proteins (Figure 2A).

**Figure 2.**
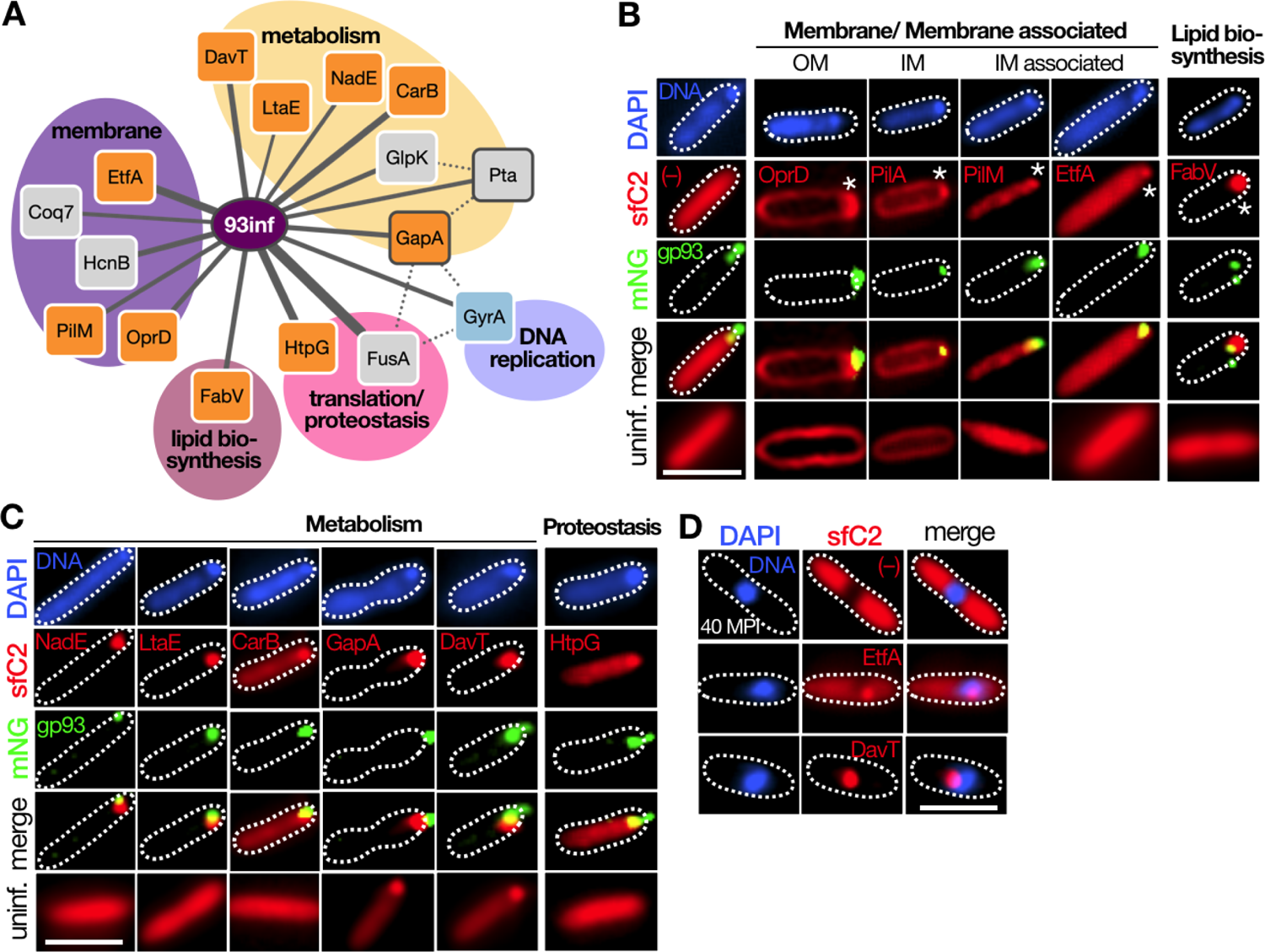
The injected protein cargo of ΦKZ interacts with host metabolic, chaperone and membrane proteins. **(A)** Interactome map of ΦKZ proteins that are detected in AP-MS with BFDR < 0.05 in 93inf samples as compared to mNG-3xF and 93cyto negative control. Colors: orange = host proteins that co-localize with injected gp93-mNG, Blue = host proteins that do not co-localize with injected gp93-mNG, Grey = microscopy not conducted; Grey border = that PPI was also found at high confidence (BFDR <= 0.05) with phage injected gp95. Line width is proportional to the average spectral count of the interacting protein (wider width = higher spectral count). Line style = solid line are interactions derived from gp93 AP-MS experiments. Dotted lines are prey-prey connections derived from the STRING database (all types of connections with STRING score >= 0.5). Representative fluorescence microscopy images of PAO1 cells expressing C-terminal sfCherry2 (sfC2) fusions of host proteins associated with **(B)** bacterial membrane/ membrane biogenesis functions or **(C)** metabolism or proteostasis functions, infected with ΦKZ: gp93-mNeonGreen (mNG) for 10 minutes **(D)** Representative fluorescence microscopy images of PAO1 cells expressing C-terminal sfCherry2 fusions of host proteins that localize to the phage nucleus at 40 minutes post infection (MPI). **(B-D)** Cells stained with DAPI to visualize DNA. Scale bars = 5 µM. See also **Supplementary Table S2**.

Prior work has demonstrated that the ΦKZ virion consists of up to ∼80 different virion proteins^19,20^. Of this list, three virion phage proteins gp92, gp97 and gp174 (Figure 1D, oval) appeared as gp93 interactors during infection (but not in gp93cyto) and are likely co-injected with gp93 and phage gDNA. To confirm that these proteins are packaged into the phage virion, we generated C-terminal sfCherry2 fusions and evaluated their colocalization with mNeonGreen-gp93 tagged endogenously in the phage genome (ΦKZ::gp93-mNG)^21^. All three proteins were indeed packaged into the virion and injected into the cell along with gp93 (Figure 1E). Both gp97 and gp92 showed high colocalization with gp93, while gp174 showed only partial colocalization (Figure 1E). These data provide validation for the AP-MS approach yielding proteins that are co-injected with gp93 and the phage DNA. However, we did not identify gp93 interactions with paralogs present in the virion at low copy^19^ (gp94 and gp163), perhaps due to their abundance. Lastly, most of the phage proteins that interact with gp93 also appeared to interact with injected gp95, when the same AP-MS experiment was conducted (Figure 1D, grey outline).

Many gp93 interactors are *Pseudomonas* host proteins associated with metabolism and bacterial membranes/lipids (Figure 2A). To investigate the possible recruitment of these proteins to the injected DNA and protein, we fused a a C-terminal sfCherry2 to the host proteins and evaluated their localization in uninfected cells and cells infected for 10 min with ΦKZ::gp93-mNG (**Supplementary Table S2**). We observed a clear relocalization of several cytosolic and membrane proteins to the point of infection (Figure 2B**, C**). Using a similar fluorescence microscopy approach as described above, we observed recruitment of host proteins that are found either within or associated with the bacterial membrane (OprD, PilM, EtfA, PilA) to the injected phage gDNA and gp93-mNG (Figure 2B) along with a protein involved in fatty acid biosynthesis, FabV (Figure 2B).

Interestingly EtfA and DavT stayed affiliated with the mature phage nucleus later in the infection, at 40 MPI (Figure 2D). With respect to these host proteins, injected gp95 only showed an interaction with Pta and GapA but not other gp93 interactors (Figure 2A, grey outline) likely because gp95 appears to dissociate from the phage gDNA-protein complex early in the phage infection cycle (**Supplementary Figure S1A**). Together, these data reveal membrane proteins associating with injected phage protein tightly associated with phage DNA and suggest that membrane recruitment or remodeling may be enacted early during phage infection.

### The injected phage genome and protein cargo of ΦKZ is associated with a lipid rearrangement

Given the high number of cell envelope and membrane proteins being recruited to the gp93-phage DNA complex, we next sought to directly investigate if indeed the early part of the infection cycle is accompanied by the rearrangement of membrane lipids to the injected phage gDNA and protein cargo. To visualize the potential recruitment of bacterial membrane lipids to the injected phage gDNA, we used fluorescence microscopy to track the localization of lipids using the lipophilic dye FM4-64 and the phage and host gDNA using DAPI staining. FM4-64 is a dye that has been used to label endocytic compartments in eukaryotic cells^22^. In phage infections arrested with gentamicin we observed that the injected phage gDNA localizes as intense DAPI-stained puncta just inside the cell envelope (Figure 3A, blue). Interestingly, with FM4-64 staining, in addition to the expected labeling of the bacterial membrane, we observed intense punctate staining colocalized with the injected ΦKZ gDNA (Figure 3A, red & overlay). We observed similar FM4-64 stained puncta colocalized with the injected phage DNA in infections of another nucleus-forming jumbo phage ΦPA3 (Figure 3A). Moreover, ΦKZ infections arrested with Cas13a targeting transcripts encoding the major nucleus protein gp54 (Figure 3B) revealed a similar staining pattern suggesting that lipid re-arrangements precede nucleus formation. FM4-64-stained puncta are not observed in samples of uninfected cells (Figure 3A**, B**), nor do DNA-containing ΦKZ or ΦPA3 virions in a cell lysate stain positive for membrane (Figure 3C).

**Figure 3.**
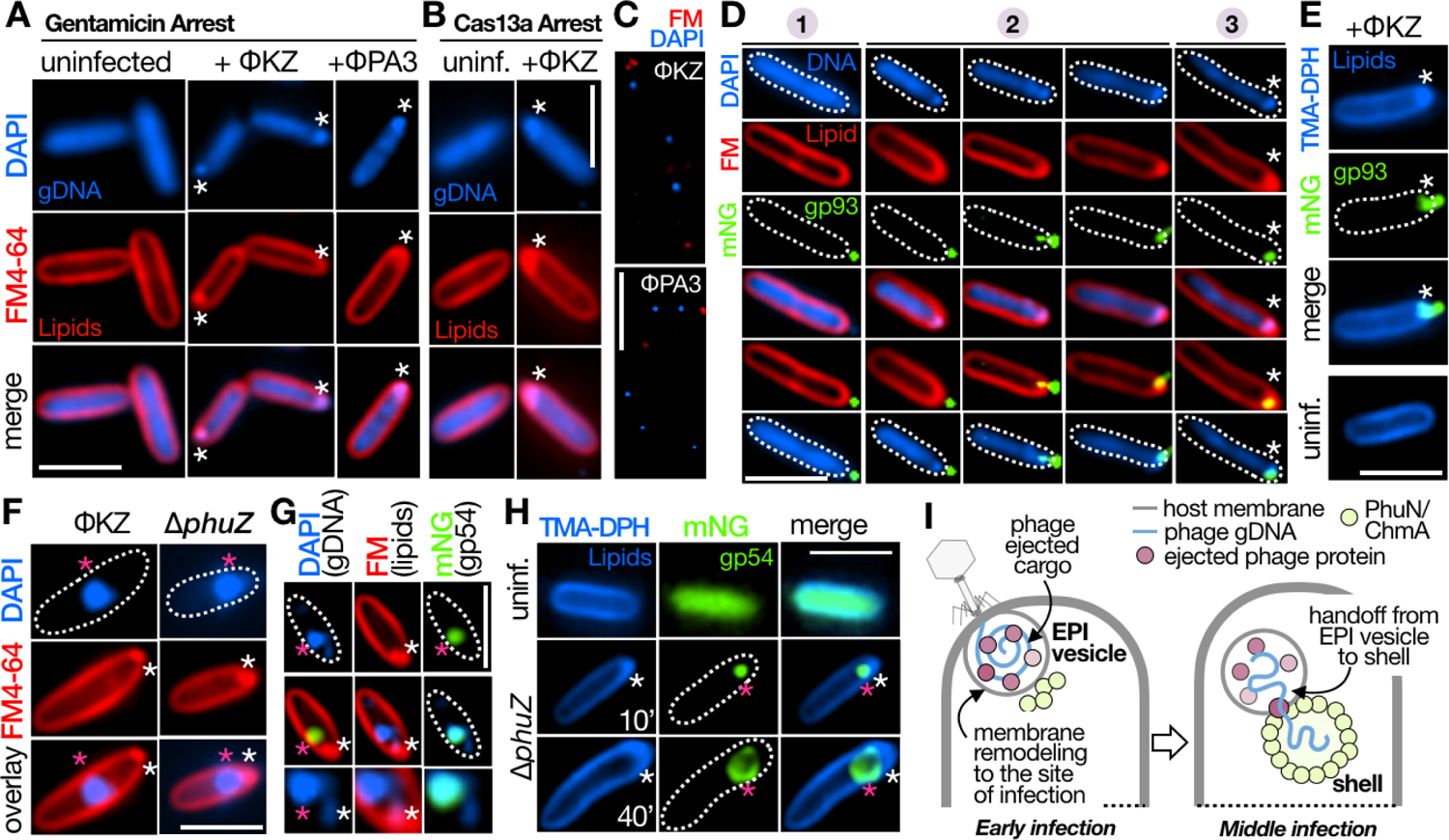
The injected gDNA and protein cargo of ΦKZ-like jumbophages associates with bacterial membrane derived lipids to produce a lipidic compartment distinct from the phage nucleus. **(A)** Representative fluorescence microscopy images of PAO1 cells: **(A)** treated with gentamicin (50 µg/mL) uninfected/ infected with ΦKZ/ ΦPA3 or, **(B)** expressing Cas13a and crRNA targeting the transcript of the major ΦKZ nucleus protein gp54; uninfected/ infected with ΦKZ. (**C**) Representative fluorescence microscopy images of ΦKZ/ ΦPA3 virions isolated from bacterial cell lysate stained with DAPI and FM4-64. (**D**) Representative fluorescence microscopy images of PAO1 cells treated with gentamicin (50 µg/mL) and infected with ΦKZ: gp93-mNeonGreen (mNG). Numbers indicate sequential stages of gp93-mNG and gDNA cargo ejection from phage, or (**E**) uninfected/ infected with ΦKZ: gp93-mNG. (**F**) Representative fluorescence microscopy images of PAO1 cells infected with WT ΦKZ/ PhuZ KO ΦKZ for 40 minutes. (**G**) Representative fluorescence microscopy images of PAO1 cells expressing mNG-gp54 infected with PhuZ KO ΦKZ for 25 minutes. **(H)** Representative fluorescence microscopy images of PAO1 cells expressing mNG-gp54 infected with PhuZ KO ΦKZ for 10 minutes or 40 minutes. **(A-B, D, F-G)** Cells or **(C)** virion samples stained with DAPI to visualize DNA and FM4-64 to visualize lipids. **(E, H**) Cells stained with TMA-DPH to visualize lipids. EPI vesicle is indicated with a white asterix. Phage nucleus is indicated with a pink asterix. Scale bars = 5 µM. gDNA: genomic DNA, mNG: mNeonGreen, FM: FM4-64, uninf: uninfected cells. **(I)** Model for the EPI vesicle compartment and its relationship to the phage nucleus.

We next tracked the localization of the phage injected protein cargo with respect to the injected gDNA and FM4-64 stained polar puncta. In gentamicin arrested infections of ΦKZ::gp93-mNG, we observed that the gp93-mNG packaged in the phage virion is co-injected with the phage gDNA and colocalizes with the FM4-64 stained puncta just inside the cell (Figure 3D). Notably, whilst observing sequential stages of phage cargo ejection (Figure 3D**, 1-3)** we observed that the FM4-64 staining was most prominent when both the phage gDNA and gp93-mNG had been completely injected into the bacteria (Figure 3D**, 3)**. To further corroborate our observation of membrane reorganization in infected cells, we additionally used the membrane permeant lipophilic dye (TMA-DPH). Using this dye, we observed a similar pattern of accumulation of membrane stain (blue) at the site of injection co-localized with the injected gp93-mNG (Figure 3E). Overall these results suggest that bacterial membranes are recruited to the injected phage gDNA and protein cargo of ΦKZ-like jumbophages during the early parts of the phage infection cycle. Based on the collective evidence from fluorescence microscopy experiments and previous cryo-ET^6,15^ we suggest that the injected phage gDNA and protein cargo associate with lipids derived from the bacterial inner membrane to assemble a small spherical compartment. To be consistent with previous work^15,23^, we refer to this body as the EPI (early phage infection) vesicle.

### The early phage infection vesicle is distinct from the phage nucleus

While the phage nucleus is bound by protein^5,6^ and not by membrane^1^, it is unclear what the spatiotemporal relationship is between the phage DNA, the EPI vesicle, and the proteinaceous phage nucleus. To address this, infections were allowed to progress for 40 min, during which the phage nucleus would assemble and the DAPI stained phage DNA would move to the middle of the cell due to the tubulin-like protein PhuZ. The FM4-64 stained puncta largely disappears at later time points, but in a small percentage of cells, the FM4-64 stained puncta were observed, where they remained at the cell pole (Figure 3F). Cells were next infected with a Δ*phuZ* mutant ΦKZ phage^21^, which limits the movement of the phage nucleus away from the cell pole, but otherwise progresses through infection normally. In this mutant, we observed that the FM4-64 stained puncta can be clearly visualized just inside the cell envelope, distinct from the DAPI stained phage nucleus (Figure 3F). To track the localization of the phage nucleus itself, we fused mNeonGreen to PhuN/Chimallin (mNG-gp54). In PAO1 cells expressing mNG-gp54 (green), infected with ΦKZ *ΔphuZ*, we observed a phage nucleus that has acquired DAPI-stained ΦKZ gDNA (blue) from the FM4-64 stained compartments (red) (Figure 3G). We orthogonally confirmed that the EPI vesicle and phage nucleus form spatially separate compartments by infecting PAO1 cells expressing mNG-gp54 with ΦKZ (PhuZ KO) and staining with lipophilic dye TMA-DPH (Figure 3H). Overall, we propose that the EPI vesicle is independent from the gp54-defined proteinaceous nucleus, which proximally receives the phage DNA in a protected manner in the middle stages of the infection once the nascent phage nucleus has formed (Figure 3I).

### The EPI vesicle is a hub for early phage transcription and translation but not DNA replication

For the EPI vesicle to be a productive step in early phage biology, we reasoned that early parts of the ΦKZ infection cycle must be carried out here. These biological processes would include immediate protection from nucleases, phage RNA transcription, mRNA export into the cytoplasm, and protein translation, prior to the DNA hand off discussed above. These steps of early phage infection will be queried below.

Early phage RNA transcription is kick-started immediately after infection by the injected virion RNA polymerase complex (vRNAP; constituted by gp80, 149, 178, and 180^9,24^). To assess whether the vRNAP is injected with the phage genome and co-localizes with the EPI vesicle, we packaged ΦKZ with C-terminal sfCherry2 fusions of vRNAP proteins gp149 or gp180. Upon infecting PAO1 cells with these phages, we observed that indeed these proteins are co-injected with the phage gDNA (Figure 4A). Interestingly at 40 MPI, we observed that although the replicating phage gDNA appeared in the mature nucleus at the mid-cell position, the vRNAP proteins continued to be observed at the cell pole (Figure 4A**, Supplementary Figure S3A**). This pattern is similar to vRNAP localization recently also shown to be injected but subsequently left behind as the DNA moves to the nucleus^12^, but in stark contrast to the localization of the injected inner body proteins (**Supplementary Figure S1A**), which move with the phage nucleus. These distinct localization patterns for injected vRNAP and IB proteins could explain why we did not observe vRNAP interacting with gp93 via AP-MS. Additionally, the vRNAP is present in low abundance, estimated at ∼10 copies per virion^12^. From these observations, we conclude that the injected phage vRNAP initiates early transcription in the EPI vesicle and is stripped off the phage genome during a regulated transfer of gDNA from the EPI vesicle to the proteinaceous nucleus.

**Figure 4.**
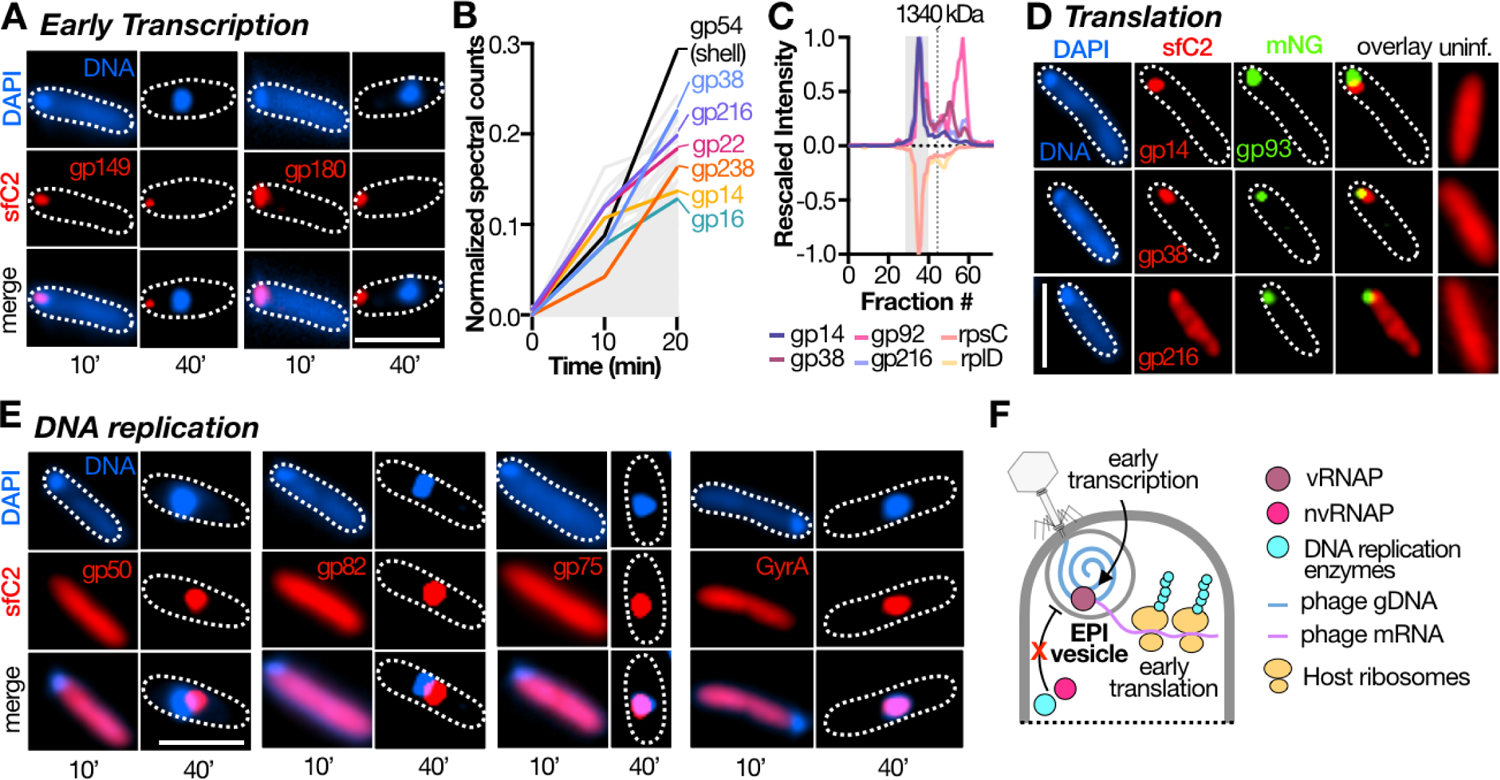
Multiple phage early-life cycle functions are carried out at the EPI vesicle. **(A)** Representative fluorescence microscopy images of PAO1 cells infected for 10 or 40 MPI with ΦKZ packaged with C-terminal sfCherry2 fusions of ΦKZ vRNAP subunits. **(B)** Line Plot of normalized spectral counts (y-axis) vs time (x-axis) of all phage proteins in PAO1 cells infected with ΦKZ. Abundantly synthesized phage proteins detected in gp93-3xFLAG AP-MS are highlighted. Spectral counts of gp54 (major nucleus protein of ΦKZ) shown for comparison. **(C)** Elution profiles of PAO1 ribosomal proteins and ΦKZ proteins that co-elute with the ribosome as determined by SEC-MS^20^. Major peak corresponding to 70S ribosomal assembly highlighted with gray rectangle. **(D)** Representative fluorescence microscopy images of PAO1 cells expressing C-terminal sfCherry2 (sfC2) fusions of ribosome associating phage proteins (gp14, 38, 216) uninfected/ infected with ΦKZ:gp93-mNG. **(E)** Representative fluorescence microscopy images of PAO1 cells expressing C-terminal sfCherry2 (sfC2) fusions of phage proteins involved with DNA polymerase/ helicase functions or PAO1 type II topoisomerase GyrA, uninfected/ infected with ΦKZ for 10 or 40 MPI. **(F)** Model for the phage life-cycle functions carried out within the EPI vesicle. **(A, D-E)** Scale bar = 5µm, mNG: mNeonGreen; sfC2: sfCherry2; uninf: uninfected cells. See also **Supplementary Figure S3** and **Supplementary Figure S4**.

After rapid phage transcription, translation of phage proteins begins. Despite the effective gentamicin arrest (i.e. the phage infection did not progress), some non-virion early expressed phage proteins (gp14, 16, 22, 38, 216, 238, 280) were also detected as interactors of gp93 (Figure 1D). We and others have previously not detected these proteins in the virion^19,20^, and thus we suggest that they are highly expressed and thus still synthesized during gentamicin treatment. Consistent with this, during unarrested infection, these proteins are among the earliest and most highly expressed, at levels comparable to the most abundantly expressed phage protein gp54, the major nucleus protein (Figure 4B and **Supplementary Table S1**). A subset of the rapidly synthesized phage proteins (namely gp014, gp038, gp216) and one injected protein (gp92, Figure 1E) enriched in the gp93inf-MS also associate with the ribosome^20,25^ (Figure 4C). Using fluorescence microscopy, we evaluated the subcellular localization of C-terminal sfCherry2 fusions of the synthesized proteins gp14, gp38, gp216 expressed in PAO1 cells (Figure 4D). While these proteins remain diffuse in their localization in uninfected cells, they are rapidly recruited to the site of infection within 10 MPI (Figure 4D). The observation of ribosome associated phage proteins interacting with injected gp93 and localizing to the EPI vesicle suggests the recruitment of ribosomes to execute localized translation. This is consistent with a cryo-ET observation that proposed polysome accumulation adjacent to the *E. coli* phage Goslar EPI vesicle^15^. A pertinent question that remains unclear is how the phage mRNA is exported from the lipid enclosure of the EPI vesicle to the cytosol in order to access ribosomes. The remaining rapidly synthesized proteins that associate with gp93 via AP-MS (gp16, 22, 238, 280) are of unknown function but gp16 and gp280 appear to be recruited to the site of infection while gp22 and gp238 form polar puncta prior to phage infection (**Supplementary Figure S4**). Overall, our gp93 AP-MS and previous SEC-MS together with data presented here suggests that specific phage proteins associate with both the EPI vesicle and host ribosomes, to potentially facilitate localized translation.

Lastly, we examined proteins that are expected to engage with phage DNA. DNA polymerases/helicases have not been identified in virion proteomes^19,20^. Therefore, the proteins needed for phage DNA replication are expected to be produced *de novo* in the cytosol and must be internalized into either the EPI vesicle or the phage nucleus. Indeed DNA replication for ΦKZ starts ∼20 min post infection^13^. We therefore tracked the subcellular localization of sfCherry2-fusions of phage-encoded DNA replication proteins (DNA polymerase: gp50, gp82; DNA Helicase: gp75) expressed in PAO1 cells with respect to the phage gDNA that is injected (10 MPI) or replicating in the nucleus (40 MPI) (Figure 4E). Unlike the host and phage proteins that are rapidly recruited to the EPI vesicle, when examining infected cells at 10 MPI, we observe that all of these proteins remain diffuse cytosolic (Figure 4E). Similarly, four different immune system proteins (HsdR, EcoRI, Cas3, Cas8) are also excluded from the EPI vesicle, similarly to the sfCherry alone negative control (**Supplementary Figure S2A**). These data collectively support previous hypotheses that phage DNA is not exposed early in the infection^3^. Later in the infection, when imaged at 40 MPI, the phage encoded (gp50, 75, 82) and host-encoded GyrA (selected because it is a gp93 interactor) DNA replication proteins are recruited to be inside or adjacent to the phage nucleus (Figure 4E). At this approximate time point is where host TopA has previously been shown to be localized inside the nucleus^1,3^ while immune enzymes are excluded ^3^. GyrA is the second host protein, together with TopA, known to localize into the phage nucleus. Our observations indicate that phage gDNA replication is compartmentalized to the phage nucleus, not the EPI vesicle. Collectively, our results establish the EPI vesicle as a hub for early phage transcription and local translation, but not DNA replication (Figure 4F).

## Discussion

The selectively permeable nucleus-like compartment assembled by ΦKZ-like jumbophages to resist DNA-targeting bacterial immune systems represents the most potent anti-immune mechanism discovered to date^3,4^. However, this structure is not built until ∼20-30 minutes post infection, and the molecular mechanisms that endow a broad spectrum of protection^3^ in early infection have remained elusive. Herein, we report that in the early stages of infection of ΦKZ, a phage-directed early phage infection (EPI) vesicle made of bacterial membrane and host proteins is assembled together with injected phage DNA and proteins. The function of this body is to coordinate early transcription and translation processes, while protecting the genome from host nucleases.

Our proteomic investigations employed an abundant injected virion protein gp93 to query the environment of the phage gDNA. Using this proteomic handle, we observed several host proteins associated with bacterial membranes, membrane biogenesis and lipid metabolism. Subsequent investigations with two lipophilic dyes revealed a strong accumulation of lipid staining at the site of infection consistent with bacterial membrane remodeling. This observation supports independent reports using cryo-ET demonstrating small circular bilayers during infection with diverse jumbo phages^6,15,23^, but having mostly unknown constituents. These observations collectively suggest a phage driven remodeling of the bacterial membrane to generate an EPI vesicle encapsulating the injected gDNA and protein cargo of ΦKZ-like jumbo phages. Notably this macromolecular compartment is entirely distinct from the phage nucleus in terms of their enclosures (membrane vs protein) and their function (i.e. early transcription in the EPI vesicle and DNA replication and middle/late transcription in the protein nucleus). Collectively, the two compartments isolate the ejected and replicating phage gDNA from immune nucleases throughout the infection cycle. Our proteomic investigations also identify multiple ribosome-associated phage proteins that may help facilitate mRNA export and localized translation at the EPI vesicle. The mechanism of mRNA and DNA export from the EPI vesicle remains a significant open question along with the roles of the many injected protein paralogs (i.e. gp92, gp93, gp94, etc. **Supplementary Figure S1**) and recruited host proteins.

Taken together, our studies provide the first insights into the molecular ‘parts-list’ of components that constitute the EPI vesicle, observed in infections of ΦKZ and other related jumbophages^6,15,23^ and help explain the diverse functions mediated by this compartment. The collective observation that this phage is creating a vesicle analogous to what follows receptor-mediated endocytosis in eukaryotic viruses represents a whole new paradigm in mechanisms employed by bacteriophages to defend against bacterial immune systems in the early parts of the infection cycle where they are most vulnerable to attack.

## Acknowledgements

This work was supported by an NIH grant 1R01AI167412 (J.B.D. and D.L.S.) and 1R01AI171041 (J.B.D. and D.A.A.). D.M. is supported by the NIH Ruth L. Kirschstein National Research Service (NRSA) Award 1F32GM149125-01.

## Additional Notes

Note that microscopy data appearing in Supplementary Figure S1A previously appeared in a preprint (https://doi.org/10.1101/2022.09.17.508391) from our lab. These data have been removed from the Li *et al*. 2022 manuscript and used here instead. The Li manuscript has been updated and these data will not appear in the final published paper.

## STAR Methods

### KEY RESOURCES TABLE

| REAGENT/ RESOURCE | SOURCE | IDENTIFIER |
| --- | --- | --- |
| **Bacterial strains** |  |  |
| *Pseudomonas aeruginosa* strain PAO1 | Joe Bondy-Denomy Lab | NCBI: NC_002516.2 |
| PAO1 [pHERD30T-PHIKZ089-mNeonGreen] | This study | PHIKZ089: Uniprot ID: Q8SD73 |
| PAO1 [pHERD30T-PHIKZ090-mNeonGreen] | This study | PHIKZ090: Uniprot ID: Q8SD72 |
| PAO1 [pHERD30T-PHIKZ093-mNeonGreen] | Guan et. al.<sup>21</sup> | PHIKZ093: Uniprot ID: Q8SD69 |
| PAO1 [pHERD30T-PHIKZ095-mNeonGreen] | This study | PHIKZ095: Uniprot ID: Q8SD67 |
| PAO1 [pHERD30T-PHIKZ097-mNeonGreen] | This study | PHIKZ097: Uniprot ID: Q8SD65 |
| PAO1 [pHERD30T-PHIKZ162-mNeonGreen] | This study | PHIKZ162: Uniprot ID: Q8SD00 |
| PAO1 [pHERD30T-PHIKZ093-3xFLAG] | Fossati et. al.<sup>20</sup> | PHIKZ093: Uniprot ID: Q8SD69 |
| PAO1 [pHERD30T-PHIKZ093C (PHIKZ093 D157-K435)-3xFLAG] | This study | PHIKZ093: Uniprot ID: Q8SD69 |
| PAO1 [pHERD30T-mNeonGreen-3xFLAG] | This study | N/A |
| PAO1 [pHERD30T-PHIKZ014-sfCherry2] | This study | PHIKZ014: Uniprot ID: Q8SDE8 |
| PAO1 [pHERD30T-PHIKZ016-sfCherry2] | This study | PHIKZ016: Uniprot ID: Q8SDE6 |
| PAO1 [pHERD30T-PHIKZ022-sfCherry2] | This study | PHIKZ022: Uniprot ID: Q8SDE0 |
| PAO1 [pHERD30T-PHIKZ038-sfCherry2] | This study | PHIKZ038: Uniprot ID: Q8SDC4 |
| PAO1 [pHERD30T-PHIKZ092-sfCherry2] | This study | PHIKZ092: Uniprot ID: Q8SD70 |
| PAO1 [pHERD30T-PHIKZ097-sfCherry2] | This study | PHIKZ097: Uniprot ID: Q8SD65 |
| PAO1 [pHERD30T-PHIKZ174-sfCherry2] | This study | PHIKZ174: Uniprot ID: Q8SCY8 |
| PAO1 [pHERD30T-PHIKZ216-sfCherry2] | This study | PHIKZ216: Uniprot ID: Q8SCU6 |
| PAO1 [pHERD30T-PHIKZ238-sfCherry2] | This study | PHIKZ238: Uniprot ID: Q8SCS4 |
| PAO1 [pHERD30T-PHIKZ238.2-sfCherry2] | This study | PHIKZ238.2: Uniprot ID: L7T0M4 |
| PAO1 [pHERD30T-PHIKZ260-sfCherry2] | This study | PHIKZ260: Uniprot ID: Q8SCQ2 |
| PAO1 [pHERD30T-PHIKZ280-sfCherry2] | This study | PHIKZ280: Uniprot ID: Q8SCN2 |
| PAO1 [pHERD30T-DAVT_PSEAE-sfCherry2] | This study | DAVT: Uniprot ID: Q9I6M4 |
| PAO1 [pHERD30T-LTAE_PSEAE-sfCherry2] | This study | LTAE: Uniprot ID: Q9HTF1 |
| PAO1 [pHERD30T-NADE_PSEAE-sfCherry2] | This study | NADE: Uniprot ID: Q9HUP3 |
| PAO1 [pHERD30T-CARB_PSEAE-sfCherry2] | This study | CARB: Uniprot ID: P38100 |
| PAO1 [pHERD30T-GAPA_PSEAE-sfCherry2] | This study | GAPA: Uniprot ID: P27726 |
| PAO1 [pHERD30T-GYRA_PSEAE-sfCherry2] | This study | GYRA: Uniprot ID: P48372 |
| PAO1 [pHERD30T-HTPG_PSEAE-sfCherry2] | This study | HTPG: Uniprot ID: Q9I3C5 |
| PAO1 [pHERD30T-FABV_PSEAE-sfCherry2] | This study | FABV: Uniprot ID: Q9HZP8 |
| PAO1 [pHERD30T-PORD_PSEAE-sfCherry2] | This study | PORD: Uniprot ID: P32722 |
| PAO1 [pHERD30T-PILM_PSEAE-sfCherry2] | This study | PILM: Uniprot ID: G3XD28 |
| PAO1 [pHERD30T-ETFA_PSEAE-sfCherry2] | This study | ETFA: Uniprot ID: Q9HZP7 |
| PAO1 [pHERD30T-PILA_PSEAE-sfCherry2] | This study | PILA: Uniprot ID: P04739 |
| PAO1 [pHERD30T-sfCherry2] | Mendoza et. al.<sup>3</sup> | SDM222 |
| PAO1 [pHERD30T-mNeonGreen-PHIKZ054] | This study | PHIKZ054: Uniprot ID: Q8SDA8 |
| PAO1 [pHERD30T-PHIKZ149-sfCherry2] | This study | PHIKZ149: Uniprot ID: Q8SD13 |
| PAO1 [pHERD30T-PHIKZ180-sfCherry2] | This study | PHIKZ180: Uniprot ID: Q8SCY2 |
| PAO1 [pHERD30T-PHIKZ050-sfCherry2] | This study | PHIKZ050: Uniprot ID: Q8SDB2 |
| PAO1 [pHERD30T-PHIKZ082-sfCherry2] | This study | PHIKZ082: Uniprot ID: Q8SD80 |
| PAO1 [pHERD30T-PHIKZ075-sfCherry2] | This study | PHIKZ075: Uniprot ID: Q8SD87 |
| PAO1 [pHERD30T-sfCherry2-HsdR] | Mendoza et. al.<sup>3</sup> | SDM223 |
| PAO1 [pHERD30T-EcoRI-sfCherry2] | Mendoza et. al.<sup>3</sup> | SDM248 |
| PAO1 [pHERD30T-sfCherry2-Cas3] | Mendoza et. al.<sup>3</sup> | N/A |
| PAO1 [pHERD30T-sfCherry2-Cas8] | Mendoza et. al.<sup>3</sup> | N/A |
| **Bacteriophages** |  |  |
| *P. aeruginosa* bacteriophage ΦKZ | Alan Davidson Lab<sup>16</sup> | NCBI: NC_004629.1 |
| *P.aeruginosa* bacteriophage ΦPA3 | David Agard Lab<sup>2</sup> | NCBI: NC_028999.1 |
| *Pseudomonas aeruginosa* bacteriophage ΦKZ:gp93-mNeonGreen (genomic fusion) | Guan et. al.<sup>21</sup> | N/A |
| ΦKZ: gp089-mNeonGreen (exogenously packaged) | This study | N/A |
| ΦKZ: gp090-mNeonGreen (exogenously packaged) | This study | N/A |
| ΦKZ: gp093-mNeonGreen (exogenously packaged) | Guan et. al.<sup>21</sup> | N/A |
| ΦKZ: gp095-mNeonGreen (exogenously packaged) | This study | N/A |
| ΦKZ: gp097-mNeonGreen (exogenously packaged) | This study | N/A |
| ΦKZ: gp162-mNeonGreen (exogenously packaged) | This study | N/A |
| ΦKZ: gp93-mNeonGreen (genomic fusion)<br>gp092-sfCherry2 (exogenously packaged) | This study | N/A |
| ΦKZ: gp93-mNeonGreen (genomic fusion)<br>gp097-sfCherry2 (exogenously packaged) | This study | N/A |
| ΦKZ: gp93-mNeonGreen (genomic fusion)<br>gp174-mNeonGreen (exogenously packaged) | This study | N/A |
| ΦKZ: gp149-sfCherry2 (exogenously packaged) | This study | N/A |
| ΦKZ: gp180-sfCherry2 (exogenously packaged) | This study | N/A |
| **Chemicals and Reagents** |  |  |
| (N-(3-Triethylammoniumpropyl)-4-(6-(4-(Diethylamino) Phenyl) Hexatrienyl) Pyridinium Dibromide (FM4-64) | ThermoFisher | Cat# T13320 |
| (1-(4-Trimethyl ammoniumphenyl)-6-Phenyl-1,3,5-Hexatriene p-Toluenesulfonate) (TMA-DPH) | ThermoFisher | Cat# T204 |
| 4',6-diamidino-2-phenylindole (DAPI) | Invitrogen | Cat# D1306 |
| Luria Broth (Miller's) | Research Products International (RPI) | Cat# L24400 |
| Magnesium Sulfate salt | RPI | Cat# M24300 |
| Gentamicin sulfate salt | Sigma-Aldrich | Cat# G1264 |
| L-(+)-Arabinose | Sigma-Aldrich | Cat# A3256 |
| Agarose, Molecular Biology Grade | RPI | Cat# A20090 |
| Trizma® base | Sigma-Aldrich | Cat# T4661 |
| Sodium Chloride salt, ACS grade | RPI | Cat# S23025 |
| EDTA Disodium Salt [Ethylenediaminetetra-acetic acid, disodium salt dihydrate] | RPI | Cat# E57020 |
| Nonidet P-40 (NP-40) | RPI | Cat# N59000 |
| cOmplete™, Mini, EDTA-free Protease Inhibitor Cocktail | Millipore-Sigma | Cat# 11836170001 |
| Benzonase | Millipore-Sigma | Cat# 101656 |
| 4x Laemmli Sample Buffer | Bio-RAD | Cat# 1610747 |
| Beta-mercaptoethanol/ 2-Mercaptoethanol | Bio-RAD | Cat# 1610710 |
| 10x Tris/Glycine/SDS Electrophoresis Buffer | Bio-RAD | Cat# 1610772 |
| PVDF Transfer Membranes, 0.2 μm | ThermoFisher | Cat# 88520 |
|  | Scientific |  |
| 10x Tris/Glycine Buffer (Western Blots, Native Gels) | Bio-RAD | Cat# 1610771 |
| Methanol ACS Grade (99.8%) | Fisher Scientific | Cat# A412-4 |
| Omniblock dry milk powder, non-fat | RPI | Cat# M17200 |
| Tween™ 20 Surfact-Amps™ Detergent Solution | ThermoFisher Scientific | Cat# PI85114 |
| Mouse anti-FLAG M2 antibody | Sigma-Aldrich | Cat# F1804 |
| Purified anti-E. coli RNA Polymerase $\beta$ Antibody | Biolegend | Cat# 663905 |
| Goat anti-mouse HRP | Invitrogen | Ref: 62-6520; Lot: XD347166 |
| Clarity Western ECL substrate | Bio-RAD | Cat# 1705061 |
| Anti-Flag M2 Magnetic Beads | Sigma-Aldrich | Cat# M8823 |
| RapiGest SF Surfactant | Waters | Cat# 186008740 |
| Urea, ACS reagent, 99.0-100.5% | Sigma-Aldrich | Cat# U5128 |
| DTT, 1,4-dithiothreitol | Sigma-Aldrich | Cat# 10197777001 |
| Iodoacetamide | Sigma-Aldrich | Cat# I6125 |
| Trypsin | Promega | Cat# VA900A |
| Trifluoroacetic acid (TFA) | Sigma-Aldrich | Cat# 8082600026 |
| Formic Acid, 99.0+%, Optima™ LC/MS Grade | Fisher Scientific | Cat# A117-05AMP |
| Acetonitrile, LC-MS Grade, 99.8%, | Fisher Scientific | Cat# AA47138K2 |
| Ammonium bicarbonate | Millipore Sigma | Cat# A6141 |
| Tris(2-carboxyethyl)phosphine hydrochloride (TCEP) | Millipore Sigma | Cat# 51805-45-9 |
| 2-Chloroacetamide | Millipore Sigma | Cat# C0267 |
| Chloroform (HPLC) | Fisher Scientific | Cat# C607-4 |
| **Critical Commercial Assays** |  |  |
| NEBuilder® HiFi DNA Assembly Master Mix | New England Biolabs (NEB) | Cat# E2621 |
| Phusion® High-Fidelity PCR Kit | NEB | Cat# E0553 |
| DNA Clean & Concentrator kit | Zymo Research | Cat #D4034 |
| Plasmid Miniprep Kit | Zymo Research | Cat #ZD4037 |
| **Deposited Data** |  |  |
| FLAG AP-MS data | This study | PXD049415 |
| PHIKZ Time Course MS data | This study | PXD050275 |
| PHIKZ SEC-MS data | Fossati 2023 | MSV000091715 |
| **Recombinant DNA** |  |  |
| pHERD30T-PHIKZ089-mNeonGreen | This study | PHIKZ089: Uniprot ID: Q8SD73 |
| pHERD30T-PHIKZ090-mNeonGreen | This study | PHIKZ090: Uniprot ID: Q8SD72 |
| pHERD30T-PHIKZ093-mNeonGreen | Guan et. al.<sup>21</sup> | PHIKZ093: Uniprot ID: Q8SD69 |
| pHERD30T-PHIKZ095-mNeonGreen | This study | PHIKZ095: Uniprot ID: Q8SD67 |
| pHERD30T-PHIKZ097-mNeonGreen | This study | PHIKZ097: Uniprot ID: Q8SD65 |
| pHERD30T-PHIKZ162-mNeonGreen | This study | PHIKZ162: Uniprot ID: Q8SD00 |
| pHERD30T-PHIKZ093-3xFLAG | Fossati et. al.<sup>20</sup> | PHIKZ093: Uniprot ID: Q8SD69 |
| pHERD30T-PHIKZ093C (PHIKZ093 D157-K435)-3xFLAG | This study | PHIKZ093: Uniprot ID: Q8SD69 |
| pHERD30T-mNeonGreen-3xFLAG | This study | N/A |
| pHERD30T-PHIKZ014-sfCherry2 | This study | PHIKZ014: Uniprot ID: Q8SDE8 |
| pHERD30T-PHIKZ016-sfCherry2 | This study | PHIKZ016: Uniprot ID: Q8SDE6 |
| pHERD30T-PHIKZ022-sfCherry2 | This study | PHIKZ022: Uniprot ID: Q8SDE0 |
| pHERD30T-PHIKZ038-sfCherry2 | This study | PHIKZ038: Uniprot ID: Q8SDC4 |
| pHERD30T-PHIKZ092-sfCherry2 | This study | PHIKZ092: Uniprot ID: Q8SD70 |
| pHERD30T-PHIKZ097-sfCherry2 | This study | PHIKZ097: Uniprot ID: Q8SD65 |
| pHERD30T-PHIKZ174-sfCherry2 | This study | PHIKZ174: Uniprot ID: Q8SCY8 |
| pHERD30T-PHIKZ216-sfCherry2 | This study | PHIKZ216: Uniprot ID: Q8SCU6 |
| pHERD30T-PHIKZ238-sfCherry2 | This study | PHIKZ238: Uniprot ID: Q8SCS4 |
| pHERD30T-PHIKZ238.2-sfCherry2 | This study | PHIKZ238.2: Uniprot ID: L7T0M4 |
| pHERD30T-PHIKZ260-sfCherry2 | This study | PHIKZ260: Uniprot ID: Q8SCQ2 |
| pHERD30T-PHIKZ280-sfCherry2 | This study | PHIKZ280: Uniprot ID: Q8SCN2 |
| pHERD30T-DAVT_PSEAE-sfCherry2 | This study | DAVT: Uniprot ID: Q9I6M4 |
| pHERD30T-LTAE_PSEAE-sfCherry2 | This study | LTAE: Uniprot ID: Q9HTF1 |
| pHERD30T-NADE_PSEAE-sfCherry2 | This study | NADE: Uniprot ID: Q9HUP3 |
| pHERD30T-CARB_PSEAE-sfCherry2 | This study | CARB: Uniprot ID: P38100 |
| pHERD30T-GAPA_PSEAE-sfCherry2 | This study | GAPA: Uniprot ID: P27726 |
| pHERD30T-GYRA_PSEAE-sfCherry2 | This study | GYRA: Uniprot ID: P48372 |
| pHERD30T-HTPG_PSEAE-sfCherry2 | This study | HTPG: Uniprot ID: Q9I3C5 |
| pHERD30T-FABV_PSEAE-sfCherry2 | This study | FABV: Uniprot ID: Q9HZP8 |
| pHERD30T-PORD_PSEAE-sfCherry2 | This study | PORD: Uniprot ID: P32722 |
| pHERD30T-PILM_PSEAE-sfCherry2 | This study | PILM: Uniprot ID: G3XD28 |
| pHERD30T-ETFA_PSEAE-sfCherry2 | This study | ETFA: Uniprot ID: Q9HZP7 |
| pHERD30T-PILA_PSEAE-sfCherry2 | This study | PILA: Uniprot ID: P04739 |
| pHERD30T-sfCherry2 | Mendoza et. al.<sup>3</sup> | pSG30T-cherry |
| pHERD30T-mNeonGreen-PHIKZ054 | This study | PHIKZ054: Uniprot ID: Q8SDA8 |
| pHERD30T-PHIKZ149-sfCherry2 | This study | PHIKZ149: Uniprot ID: Q8SD13 |
| pHERD30T-PHIKZ180-sfCherry2 | This study | PHIKZ180: Uniprot ID: Q8SCY2 |
| pHERD30T-PHIKZ050-sfCherry2 | This study | PHIKZ050: Uniprot ID: Q8SDB2 |
| pHERD30T-PHIKZ082-sfCherry2 | This study | PHIKZ082: Uniprot ID: Q8SD80 |
| pHERD30T-PHIKZ075-sfCherry2 | This study | PHIKZ075: Uniprot ID: Q8SD87 |
| pHERD30T-sfCherry2-HsdR | Mendoza et. al.<sup>3</sup> | pSDM153 |
| pHERD30T-EcoRI-sfCherry2 | Mendoza et. al.<sup>3</sup> | pSDM171 |
| pHERD30T-sfCherry2-Cas3 | Mendoza et. al.<sup>3</sup> | pSG30T-cherrycas3(IC) |
| pHERD30T-sfCherry2-Cas8 | Mendoza et. al.<sup>3</sup> | pSG30T-cherrycas8(IC) |
| **Softwares and Algorithms** |  |  |
| SAINTexpress (version 3.6.1) | Teo et. al.<sup>26</sup> | https://saint-apms.sourceforge.net/Main.html |
| Fragpipe (version 19.1) | Yu et. al.<sup>27</sup> | https://fragpipe.nesvilab.org/ |
| PhageMAP | Fossati et. al.<sup>20</sup> | https://phagemap.ucsf.edu/home/ |
| Prism version 10.1.0 (264) | GraphPad Prism | https://www.graphpad.com/ |
| Fiji (Fiji is just ImageJ) (version 2.0.0-rc-69/1.53h) | ImageJ wiki | https://imagej.net |

### Fluorescence microscopy and imaging

PAO1 cells (WT or expressing sfCherry2/ mNeonGreen fusions) were grown overnight in 3 mL LBM (LB with 10mM MgSO_4_) (supplemented with gentamicin 50 μg/ml, as appropriate) at 37 °C with aeration at 175 rpm. The overnight culture was diluted 100-fold in 3 ml of LBM (containing 0.03% Arabinose) and grown at 37 °C with 175 r.p.m. shaking to an OD600 of approximately 0.5. Thereafter PAO1 cells (≈1 OD equivalent) were infected with ΦKZ particles (approximate MOI of 1-10) on ice for 10 min (to allow complete adsorption of virions onto cells) and then incubated at 30 °C for indicated times (10 or 40 minutes). 2 µL of the uninfected/ infected culture was applied onto an was gently placed onto a piece of agarose pad (approximately 1 mm thick) with 1:4 diluted LBM, arabinose (0.8%) and fluorescent dyes namely: 4′,6-diamidino-2-phenylindole (DAPI; 5 µg ml−1; Cat# D1306; Invitrogen) or (N-(3-Triethylammoniumpropyl)-4-(6-(4-(Diethylamino) Phenyl) Hexatrienyl) Pyridinium Dibromide (FM4-64; 5 µg ml−1; Cat# T13320, ThermoFisher) or (1-(4-Trimethyl ammoniumphenyl)-6-Phenyl-1,3,5-Hexatriene p-Toluenesulfonate) (TMA-DPH; 5 µg ml−1; Cat# T204). A coverslip (no. 1.5; Thermo Fisher Scientific) was gently laid over the agarose pad and the sample was imaged under a fluorescence microscope at 30 °C within a cage incubator to maintain temperature and humidity. Microscopy was performed on an inverted epifluorescence microscope (Ti2-E; Nikon) equipped with the Perfect Focus System and a Photometrics Prime 95B 25-mm camera. Image acquisition and processing were performed with the Nikon Elements AR software v.5.02.00 (64-bit). Specimens were imaged at a time interval through channels of phase contrast (200 ms exposure for cell recognition), blue (DAPI, 200 ms exposure for phage DNA; TMA-DPH, 500 ms exposure), green (mNeonGreen, 5s long exposure for gp93-mNeonGreen) and red (FM4-64, 200ms exposure; sfCherry2, 500ms exposure). Images were analyzed with Fiji/ ImageJ (2.0.0-rc-69/1.53h)^28^

### Western Blotting

PAO1 cells (WT or expressing gp93-3xFLAG/ mNeonGreen-3xFLAG) were grown overnight in 3 mL LB (supplemented with gentamicin 50 μg/ml, as appropriate) at 37 °C with aeration at 175 rpm. Cells were diluted 1:100 from a saturated overnight culture into 5 mL LB with 10 mM MgSO4 (for WT PAO1), or supplemented with Arabinose (for PAO1: gp93-3xFLAG/ mNeonGreen-3xFLAG) and grown for 2.5 h at 37 °C with aeration at 175 rpm. Upon reaching 0.5 OD (600 nm), gentamicin was added to WT PAO1 cells (50 μg/ml), and the cells were chilled on ice for 5 min. Thereafter (1) WT PAO1 cells (≈1 OD equivalent) were infected with ΦKZ particles packaged with gp93-3xFLAG-tag/ (2) PAO1 cells expressing gp93C-3xFLAG/ mNeonGreen-3xFLAG were infected with WT ΦKZ particles (MOI ≈ 1) on ice for 10 min (to allow complete adsorption of virions onto cells) and then incubated at 30 °C for 15 min. Thereafter, the cell cultures were transferred to pre-chilled 15 mL falcon tubes and centrifuged at 6000×g, 4 °C for 5 min. The supernatant was discarded, and the cell pellet was washed twice with 2 mL of pre-chilled (4 °C) LB to remove excess unbound virions. The cell pellet was lysed in 100 μL of lysis buffer (50 mM Tris (pH 7.4), 150 mM NaCl, 1 mM EDTA, 0.5% NP-40, 1x protease inhibitor cocktail (Roche, complete mini EDTA free), 125 U Benzonase/mL). The lysed suspension was further sonicated on ice using a Q125 sonicator (10 pulses, 1 s ON, 1 s OFF, 30% amplitude). The cell lysate was centrifuged at 15,000 × g (15 min, 4 °C) to remove cellular debris. The clarified cellular lysate (100 μL) was boiled with 33 μL of 4x Laemmli Buffer (with Beta-mercaptoethanol) for 10 min. 14 μL of lysate samples were loaded. For virion control samples, 10 μL of purified virions were boiled with 3.3 μLL of 4x Laemmli Buffer (with Beta-mercaptoethanol) for 10 min, and 2 μL of samples were loaded. SDS-PAGE gels were run with running buffer (100 mL 10× Tris-Glycine SDS Buffer, 900 mL Milli-Q water) at 130 V for 1 h (constant voltage setting). The SDS-PAGE gels were transferred onto 0.2 μM PVDF membranes using a wet transfer (Transfer Buffer: 100 mL 10× Tris-Glycine Buffer, 200 mL methanol, 700 mL Milli-Q water; 100 V, 1 hour, 4 °C). The membranes were incubated with a blocking buffer (5% Omniblock milk, non-fat-dry in 1× TBST (200 mL Tris Buffer Saline, 0.20 mL Tween-20)) for 1 h at room temperature. Thereafter the blocking buffer was discarded, and the membranes were incubated with 1:1000 dilutions of mouse anti-FLAG M2 antibody (Sigma-Aldrich, Cat# F1804) or anti-RNAPᵦ antibody (Biolegend, Cat# 663905) in 1×TBST (overnight, 4 °C, with constant shaking). Thereafter the membranes were washed thrice for 10 min with TBST and incubated with 1:3000 dilution of Goat anti-mouse HRP (Ref: 62-6520; Lot: XD347166 Invitrogen) in blocking buffer for 1 hr at room temperature with constant shaking. Finally, the membranes were washed thrice for 10 min with TBST and incubated with Clarity Western ECL substrate. Membranes were imaged on an Azure 500 imager for variable amounts of time.

### FLAG AP-MS

Lysate samples of PAO1 cells prepared (prepared exactly as described for western blotting) were used for AP-MS experiments. For FLAG purifications, 30 μL of bead slurry (Anti-Flag M2 Magnetic Beads, Sigma) was washed twice with 1 mL of ice-cold wash buffer (50 mM Tris pH 7.4, 150 mM NaCl, 1 mM EDTA) and the lysate was incubated with the anti-FLAG beads at 4 °C with rotation for 2 hrs. After incubation, flow-through was removed and beads were washed once with 500 μL of wash buffer with 0.05% NP40 and twice with 1 mL of wash buffer (no NP40). Bound proteins were eluted by incubating beads with 15 μL of 100 ug/ml 3xFLAG peptide in 0.05% RapiGest in the wash buffer for 15 min at RT with shaking. Supernatants were removed and elution was repeated. Eluates were combined and 10 μL of 8 M urea, 250 mM Tris, 5 mM DTT (final concentration ∼1.7 M urea, 50 mM Tris, and 1 mM DTT) was added to give a final total volume of ∼45 μL. Samples were incubated at 60°C for 15 min and allowed to cool to room temperature. Iodoacetamide was added to a final concentration of 3 mM and incubated at room temperature for 45 min in the dark. DTT was added to a final concentration of 3 mM before adding 1 μg of sequencing-grade trypsin (Promega) and incubating at 37°C overnight. Samples were acidified to 0.5% TFA (ph<2) with 10% TFA stock and incubated for 30 min before desalting on C18 ultra micro spin columns (The Nest Group). After lyophilization of desalted eluants, samples were resuspended in 20 μL of 0.1% formic acid and 2μL were separated by a reversed-phase gradient from 2.4% acetonitrile in 0.1% formic acid to 22.4% acetonitrile over the course of 44 min, followed by a 5 min ramp to 36% acetonitrile, and a final column wash at 70.4% acetonitrile for 10 min. Separations were performed using a nanoflow 75μm ID x 25cm long picotip column packed with 1.9μM C18 particles (Dr. Maisch). Peptides were directly injected over the course of a 60 min acquisition into an Orbitrap Fusion Lumos Tribrid Mass Spectrometer (Thermo) and MS1 scans were collected in the orbitrap, while MS2 was performed with HCD fragmentation and data collection in the ion trap. Instrument performance was monitored by QCloud 2.0^29^. Raw MS data were searched against both the ΦKZ and the and the PAO1 proteomes, and using the default settings in MaxQuant (version 2.0.3.0)^30^. Peptides and proteins were filtered to 1% false discovery rate in MaxQuant, and identified proteins spectral counts were then subjected to protein-protein interaction scoring with SAINTexpress (version 3.6.1)^26^, using both the gp93cyto and the mNeongreen-3xFlag as negative controls. All raw data files and search results are available from the Pride partner ProteomeXchange repository under the PXD049415 identifier^31,32^. They can be accessed with the following login credentials: Username, Password: XFGsmnfJ.

### Time Course MS

Mid-log cultures of PAO1 cells (OD 0.5-0.6, 50 mL) were infected with WT ΦKZ particles (MOI ≈ 1) on ice for 10 min (to allow complete adsorption of virions onto cells) and then incubated at 30 °C for 10 and 20 min respectively. Cultures of infected cells were flash frozen on liquid nitrogen and subsequently mechanically lysed using a SPEX-freezer mill (cryo-milling). Powdered lysate samples were resuspended in Urea lysis buffer (8M urea, 100 mM ammonium bicarbonate pH 8) supplemented with a Complete protease inhibitor tablet (Roche). Samples were centrifuged at 13’000 rpm for 10 minutes to remove cellular debris. Protein amount was assessed by BCA and approximately ∼50 ug per sample were used per replicate. TCEP was added to 5 mM final concentration and samples were incubated at 25 C for 30 minutes on a shaker (400 rpm), reduced cysteines were alkylated with 10 mM chloroacetamide for 1 hr in the dark at 25 C. Urea concentration was reduced to 1 M by addition of 100 mM ammonium bicarbonate pH 8. 1 ug of trypsin was added per sample and proteins were digested overnight at 30 C on a thermo shaker (400 rpm).

The next day, samples were acidified with 20% TFA to pH ∼2 and desalting was performed as previously described for AP-MS. The MS acquisition was performed on an Orbitrap Exploris operating in DDA mode interfaced with a Vanquish Neo HPLC. The samples were separated in 70 minutes of a linear gradient from 5 to 37% B (0.1% FA in ACN) followed by 10 minutes at 90% B and 5 minutes to equilibrate the column at 5% B. The MS samples were searched against a protein database composed of phiKZ (refSeq id: NC_004629) and PAO1 (6026 entries in total) in Fragpipe (v19.1) using the ‘LFQ-MBR’ workflow with default parameters. All raw data files and search results from this experiment are available from the Pride partner ProteomeXchange repository under the PXD050275 identifier^31,32^. They can be accessed with the following login credentials: Username, Password: Yj0sgAHD.

### SEC-MS

SEC-MS data is derived from experiments performed for a prior publication^20^. Graphs displaying elution profiles of individual proteins were generated with Graphpad Prism.

## SUPPLEMENTARY MATERIALS

**Supplementary Figure S1.**
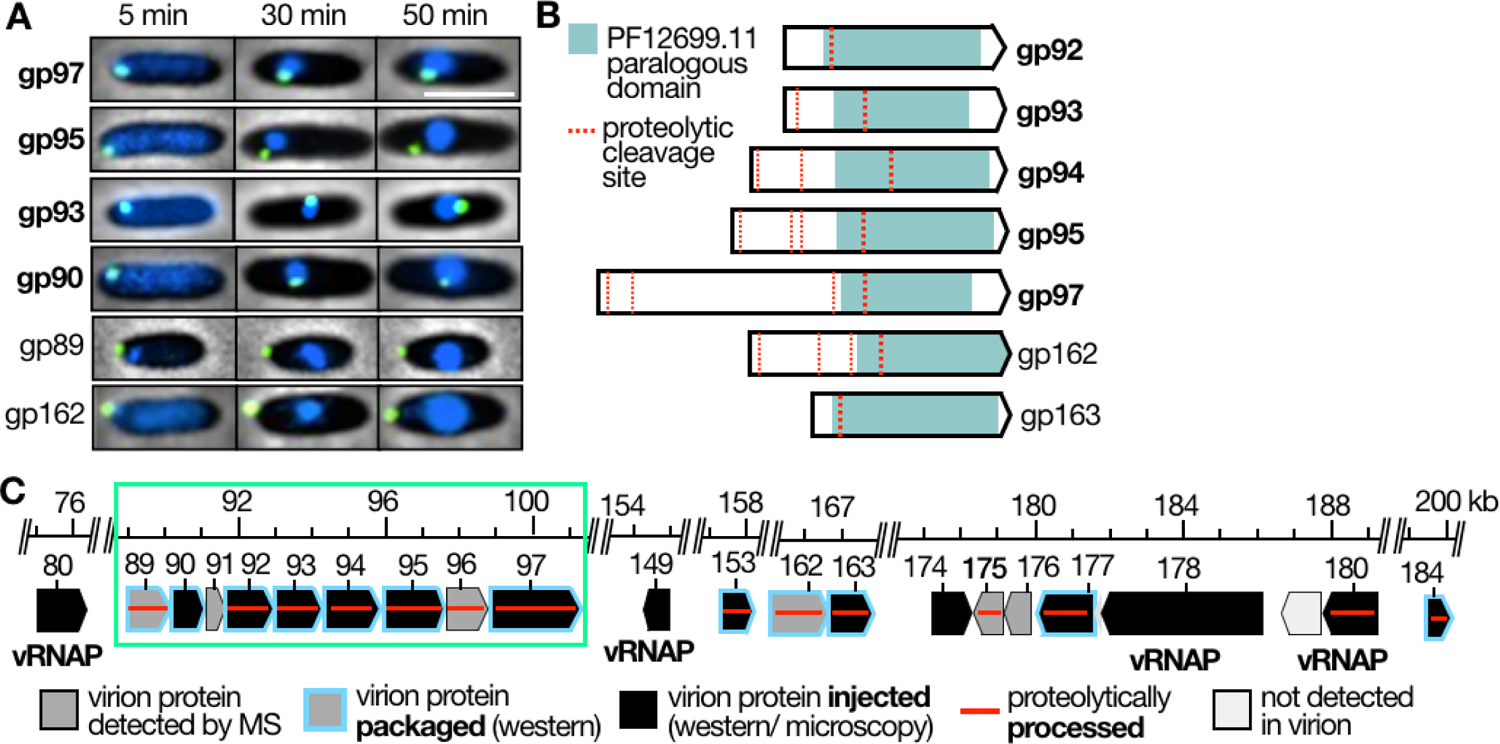
**(A)** Representative fluorescence microscopy images of PAO1 cells infected with ΦKZ packaged with C-terminal mNeonGreen (mNG) fusions of the six reported inner body (IB) proteins ^19^. Scale Bar = 5 µM **(B**) Domain diagram illustrating the paralogous domain (PF12699.11; ΦKZ-like phage internal head proteins; green) shared by gp93 and other IB and injected IB-locus proteins. Reported proteolytic cleavage sites within paralogs^19^ indicated with broken red line. (**C**) Operon diagram and genomic location of all injected ΦKZ virion proteins reported to date. IB operon highlighted with green rectangle

**Supplementary Figure S2.**
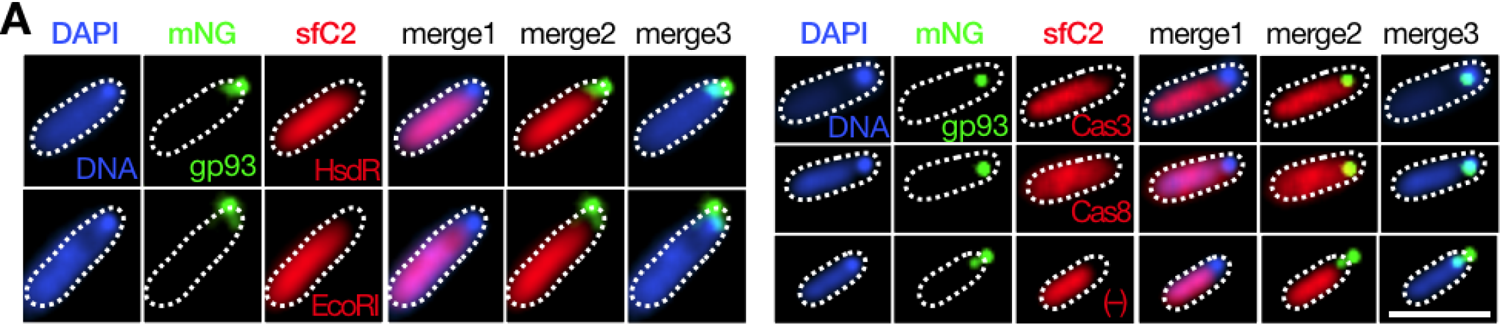
**(A)** Representative fluorescence microscopy images of PAO1 cells expressing sfCherry2 fusions of DNA-targeting immune enzymes HsdR, EcoRI, Cas3, Cas8 infected with ΦKZ: gp93-mNG for 10 minutes. Cells stained with DAPI to visualize gDNA. Scale bar = 5µm, gDNA = genomic DNA, sfC2 = sfCherry2, mNG = mNeonGreen.

**Supplementary Figure S3.**
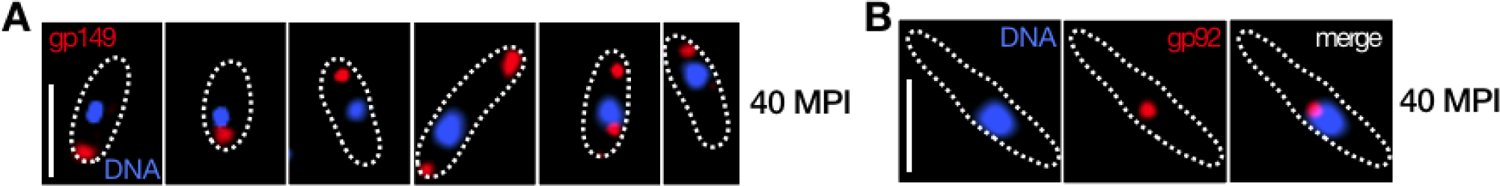
Representative fluorescence microscopy images of PAO1 cells infected with ΦKZ particles packaged with C-terminal sfCherry2 fusions of **(A)** gp149 or **(B)** gp92 imaged at 40 minutes post infection (MPI). Scale bars = 5µm, gDNA: genomic DNA, red: sfCherry2.

**Supplementary Figure S4.**
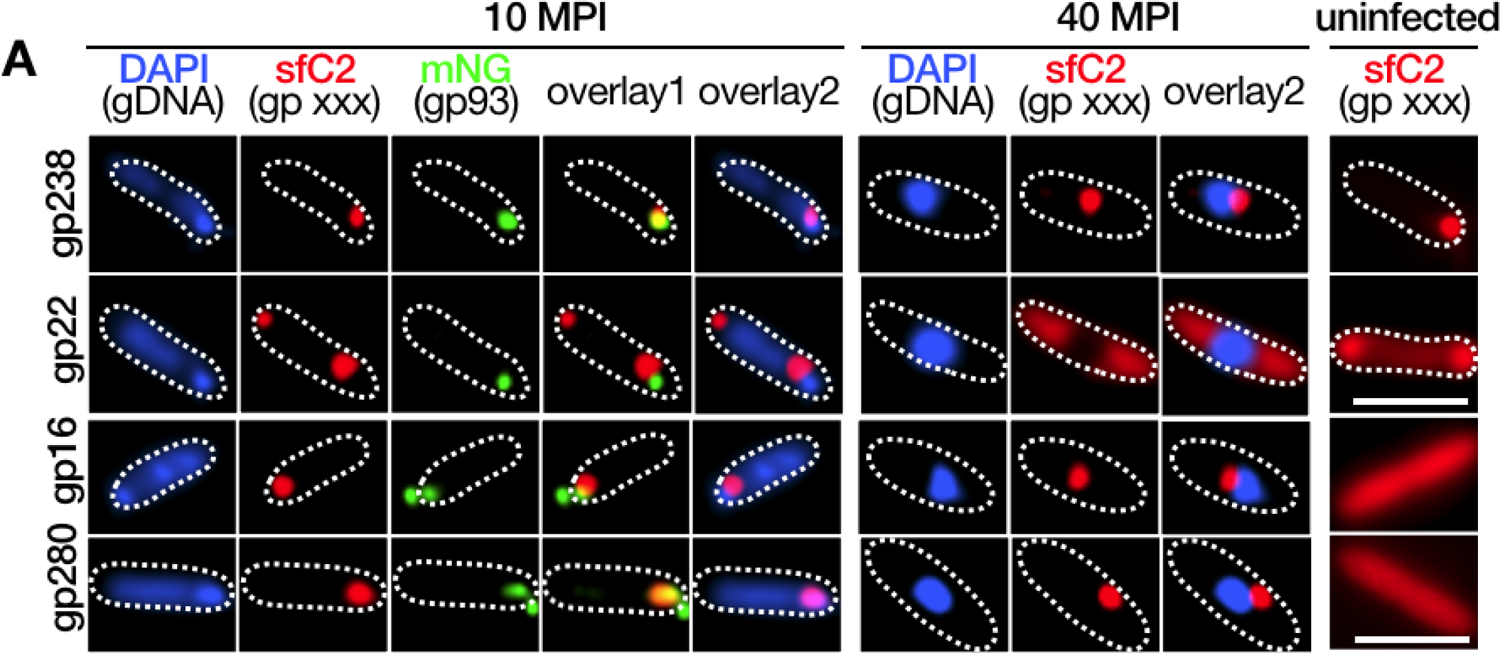
**(A)** Representative fluorescence microscopy images of PAO1 cells infected with ΦKZ particles packaged with C-terminal sfCherry2 fusions of gp238/ gp22/ gp16/ gp280 imaged at 10 and 40 minutes post infection (MPI) with ΦKZ: gp93-mNG. Cells stained with DAPI to visualize gDNA. Scale bars = 5µm, gDNA: genomic DNA, sfC2: sfCherry2, mNG: mNeonGreen.

---

**Supplementary Table S1.** Details of phage proteins detected in gp93 AP-MS. *1: ^20^,virion proteins detected in MS analysis of CsCl gradient purified phage virions; *2: ^25^,ribosome interacting protein as determined by Grad-MS; *3: ^20^,ribosome interacting protein as determined by SEC-MS. EPIV = Early Phage Infection Vesicle.

| **PHIKZ Protein** | **Virion or Synthesized** | **Localization at 10 MPI** | **Localization at 40 MPI** | **Ribosome Associated** | **Additional comments** |
| --- | --- | --- | --- | --- | --- |
| **gp014** | Synthesized | Punctate, EPIV | Diffuse, Cytosolic | Yes *2,3 |  |
| **gp016** | Synthesized | Punctate, EPIV | Punctate, Nucleus adjacent |  |  |
| **gp022** | Synthesized | Punctate | Diffuse, Cytosolic |  |  |
| **gp038** | Synthesized | Punctate, EPIV | Diffuse, Cytosolic | Yes *3 |  |
| **gp092** | Virion *1 | Punctate, EPIV | Punctate, Nucleus Adjacent | Yes *3 | IB operon protein |
| **gp097** | Virion *1 | Punctate, EPIV | Punctate, Nucleus Adjacent |  | IB operon protein |
| **gp174** | Virion *1 | Punctate, EPIV | Punctate, Polar |  |  |
| **gp189** | Synthesized | N/A | N/A |  | Not examined |
| **gp216** | Synthesized | Punctate, EPIV | Diffuse Cytosolic | Yes *3 |  |
| **gp238** | Synthesized | Punctate, EPIV | Punctate, Nucleus adjacent |  |  |
| **p72** | Synthesized | Diffuse, Cytosolic | Diffuse, Cytosolic |  |  |
| **gp266** | Synthesized | Diffuse, Cytosolic | Diffuse, Cytosolic |  | Cells display extreme filamentation |
| **gp280** | Synthesized | Punctate, EPIV | Punctate, Nucleus Adjacent |  |  |

**Supplementary Table S2.** Details of host proteins detected in gp93 AP-MS. *1: Annotation of sub-cellular localization and function as per https://www.pseudomonas.com/. EPIV = Early phage infection vesicle.

| Host Protein | PA gene number | Full protein Name | Membrane/ Cytoplasmic *1 | Membrane/ Lipid biosynthesis? *1 | Localizes to EPIV, Nucleus? |
| --- | --- | --- | --- | --- | --- |
| **LtaE** | PA5413 | Low specificity L-threonine aldolase | Unknown |  | EPIV |
| **NadE** | PA4920 | NH(3)-dependent NAD(+) synthetase | Cytoplasmic |  | EPIV |
| **PilM** | PA5044 | Type IV pilus inner membrane component PilM | Membrane | IM protein | EPIV |
| **OprD** | PA0958 | PorinD | Membrane | OM protein | EPIV |
| **GapA** | PA3195 | Glyceraldehyde-3-phosphate dehydrogenase | Cytoplasmic |  | EPIV |
| **Pta** | PA0835 | Phosphate acetyltransferase | Cytoplasmic |  | Not examined |
| **CarB** | PA4756 | Carbamoyl-phosphate synthase large chain | OMV/ Cytoplasmic |  | EPIV |
| **EtfA** | PA2951 | Electron transfer flavoprotein subunit $\alpha$ | Periplasmic | | EPIV, Nucleus adjacent (40m) |
| **HtpG** | PA1596 | Chaperone protein HtpG | OMV/ Cytoplasmic |  | EPIV |
| **FabV** | PA2950 | Enoyl-[acyl-carrier-protein] reductase [NADH] | Cytoplasmic | Fatty acid metabolism/ biosynthesis | EPIV |
| **DavT** | PA0266 | 5-aminovalerate aminotransferase DavT | Cytoplasmic |  | EPIV, Nucleus adjacent (40m) |
| **Coq7** | PA0655 | 3-demethoxyubiquinol 3-hydroxylase | Membrane | Membrane protein | Not examined |
| **HcnB** | PA2194 | Hydrogen cyanide synthase subunit HcnB | Membrane | Membrane protein | Not examined |
| **GyrA** | PA3168 | DNA gyrase subunit A | Cytoplasmic |  | Nucleus (40m) |
| **Glpk2** | PA3582 | Glycerol kinase 2 | Cytoplasmic |  | Not examined |
| **FusA** | PA4266 | Elongation factor G 1 | Cytoplasmic |  | Not examined |

